# Synthetic activation of gibberellin signaling reveals spatial coordination of root growth

**DOI:** 10.64898/2026.04.30.721855

**Authors:** Yuichiro Yagami, Ryotaro Yamada, Yuuma Ishikawa, Eiki Meguro, Kenichiro Itami, Wolf B. Frommer, Shinya Hagihara, Masayoshi Nakamura

**Author notes:** Corresponding author: Masayoshi Nakamura.

## Abstract

Gibberellins (GAs) influence cell division and elongation, profoundly shaping plant architecture and yield. GA perception occurs when bioactive GAs bind the receptor GID1, promoting DELLA degradation and activating transcriptional programs. While GA signaling in the root endodermis is essential for promoting root elongation, functions of other layers in spatial control of GA responses have not been explored. Here, we developed a synthetic GA (sGA) that does not bind endogenous GID1, together with a modified GID1 (mGID1) engineered to selectively recognize sGA, enabling cell-specific activation of GA signaling in vivo. Using this system in *Arabidopsis*, we demonstrate that coordinated action of GA signaling in the endodermis, epidermis, and other layers is required for full root elongation. Moreover, cell type-specific expression of GA biosynthetic enzymes indicates the existence of intercellular GA transport. The sGA–mGID1 system provides a versatile platform for spatially precise reprogramming of hormone signaling, enabling synthetic control of developmental processes such as root-shoot growth balance, thereby advancing applications in plant synthetic biology and sustainable crop improvement.

## Main

Gibberellins (GAs) are phytohormones involved in the regulation of a diverse range of physiological and developmental processes, including seed germination, cell proliferation and elongation, floral induction, and ovary development^1,2^. Developments during the Green Revolution demonstrated the agricultural significance of GA signaling, as mutations impairing GA biosynthesis led to reduced plant height and improved lodging resistance, thereby increasing crop yield^3–5^. This key advance and the research over the 75 years underscore the potential for enhancing crop performance provided by a higher resolution understanding of the spatiotemporal dynamics of GA signaling.

Despite nearly a century of research since the discovery of GAs, precise control of GA signaling at the cell-, tissue-, or organ-specific level remains technically challenging. The GA receptor GIBBERELLIN INSENSITIVE DWARF1 (GID1) forms a complex with GA to promote the degradation of DELLA proteins (key repressors of GA signaling), triggering downstream transcriptional reprogramming^6–10^. A classical approach to dissect cell type-specific signaling is to express the receptor in targeted cells in a receptor-deficient background. However, Arabidopsis *gid1* triple mutants exhibit severe dwarfism and sterility, rendering them genetically intractable^11^. Furthermore, the intercellular mobility of GA indicates that signaling can be activated outside GA-producing cells. Attempts to inhibit GA signaling by expressing DELLA proteins in specific cell types are also hindered by the complexity due to the existence of five distinct DELLA isoforms^12^.

Previous studies had implicated the endodermis as a critical site for GA action in root elongation^13^. GIBBERELLIC ACID INSENSITIVE (GAI), one of the DELLA proteins, inhibits root growth when a degradation-resistant mutant is expressed in the endodermis^14^. In addition, fluorescently labeled GA_3_ (GA_3_ - Fl) preferentially accumulated in endodermal cells within the root elongation zone^15^. These observations indicated that GA signaling in the endodermis is essential for root growth. Although *DELLA* genes are expressed across multiple cell layers beyond the endodermis^16,17^, their function within these layers remain unresolved. Consequently, an important question arises regarding whether GA signaling in the endodermis alone is sufficient to sustain full root growth or whether additional cell types may also contribute to this process.

Here, we developed a synthetic GA/modified GA receptor system comprising a biologically inactive synthetic GA analog (sGA) and a modified GID1 receptor (mGID1) engineered to selectively respond to sGA but not GA_3_ or GA_4_. Selective expression of mGID1 in specific cell layers and application of sGA enables reconstitution of GA signaling with high spatial and temporal resolution. By implementing the sGA-mGID system in Arabidopsis GA-deficient mutants, we demonstrated that GA signaling in both the endodermis and epidermis coordinately promotes root elongation, thereby providing direct evidence for cell-type-specific contributions to GA-mediated growth. The cell type-specific roles of GA signaling are further supported by the finding that GA signaling and the expression of the GA biosynthetic enzyme *GA 3-oxidase* (*GA3ox*) are present in different regions of the root and show only a partially overlapping pattern, in particular in the differentiation zone. The spatial separation between biosynthesis and signaling supports the concept of intercellular GA translocation, highlighting the intricate dynamics of endogenous GA.

The sGA–mGID1 system enables precise manipulation of GA signaling across multiple biological scales, ranging from individual cells and tissues to organs, subcellular compartments, and even distinct individuals within plant populations, thereby facilitating synthetic reprogramming of GA signal responses. Importantly, this approach enables the decoupling of shoot and root growth, offering a strategy to selectively enhance root development. Such targeted manipulation may mitigate the drawbacks of the Green Revolution by improving nutrient acquisition efficiency and reducing dependence on intensive fertilizer input. Collectively, these advances provide a powerful framework for dissecting hormone signaling and for optimizing plant growth and agricultural productivity with high spatial precision.

## Results

### Synthesis of sterically-hindered GA and engineering of GID1 to recognize synthetic GA

To engineer a bipartite system for chemical control over GA signaling, we synthesized GA analogs that are not recognized by native GA receptors due to steric hindrance and engineered the GA receptor to selectively recognize these analogs (Fig. 1a). Major bioactive GAs, such as GA_1_, GA_3_ and GA_4_, are C_19_-GAs characterized by a C-3β hydroxyl group, a γ-lactone ring bridging C-4 and C-10, a C-6 carboxyl group, and absence of a hydroxyl group on C-2^18,19^(Fig. 1b). These structural features are considered critical for recognition of bioactive GA by the GA-binding pocket of GID1 in the GID1 and DELLA/GAI receptor module^9,10,20^. To preserve these essential characteristics while abolishing the ability to bind to endogenous GID1, we synthesized two sGA candidates: sGA-1, in which a thiophenol was introduced to the C-1 position of GA_3_, and sGA-2, in which a benzylmercaptan was introduced to the C-1 position of GA_3_ (Fig. 1b). sGA-1 and sGA-2 had no effect on root growth, indicating that sGA-1 and sGA-2 are biologically inert toward endogenous GID1 (Extended Data Fig. 1).

**Fig. 1.**
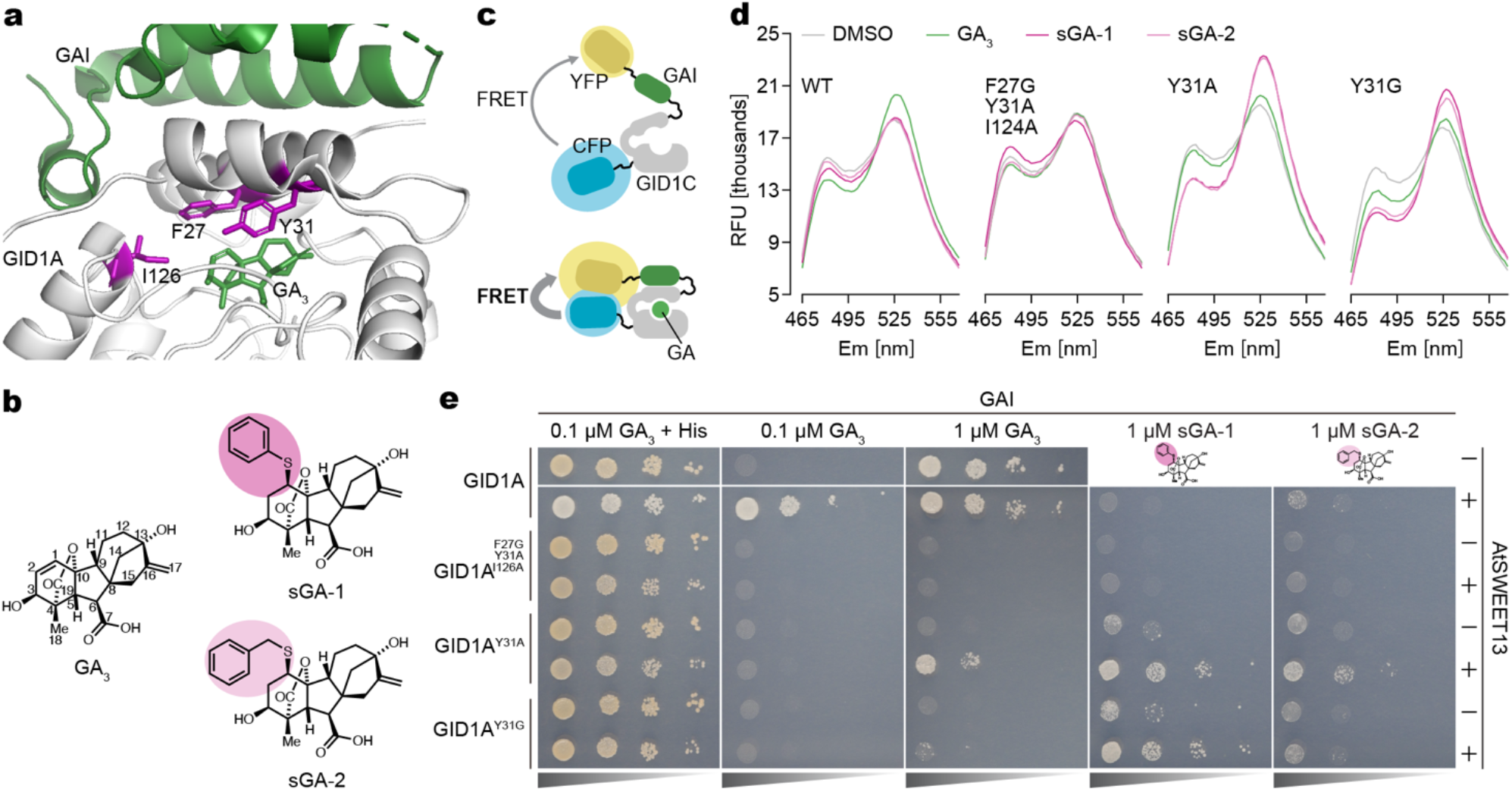
Synthesis of GA analogs and generation of GID1 that recognizes synthetic GAs. **a**, Crystal structure of GID1A (white protein) in complex with GA_3_ (small, green compound) and a truncated form of GAI (green protein). These structures are derived from a previously published ternary complex of *Arabidopsis thaliana* GID1A, GA_3_, and a GAI truncation (PDB 2ZSH). Residues in GID1A that were mutagenized to generate mGID1 candidates are highlighted in magenta. **b**, Chemical structures of GA_3_ (left, with carbon backbone numbered), sGA-1, and sGA-2. sGA-1 has a structure in which thiophenol is introduced to the C-1 position of GA_3_ (dark magenta), while sGA-2 incorporates a benzylmercaptan at the same position (light magenta). **c**, Schematic diagram of GPS1 biosensor (top). GPS1 includes GID1C as a component. When GPS1 recognizes GA, GPS1 emission ratios are altered (bottom). **d**, Fluorescence emission spectra of purified GPS1, GPS1 (GID1C^F27G, Y31A, I124A^), GPS1 (GID1C^Y31A^) or GPS1 (GID1C^Y31G^) proteins following treatment of DMSO, 1 µM GA_3_, 1 µM sGA-1, or 1 µM sGA-2. Superscripts indicate mutations introduced into GID1C, and each graph is labeled accordingly. RFU: relative fluorescence units. **e**, GA-dependent interaction assay between GID1A and GAI using yeast three-hybrid (Y3H) system. Shown are representative images. Yeast strains with or without *AtSWEET13* expression were tested. Interactions between GAI and GID1A, GID1A^F27G, Y31A, I126A^, GID1A^Y31A^, or GID1A^Y31G^ were assessed under GA_3_, sGA-1, or sGA-2 treatment. For each strain, yeast solutions of OD_600_ = 1.0, 0.1, 0.01, and 0.001 were spotted from left to right.

To develop mGID1 variants capable of recognizing the bulky sGA candidates, we employed a structure-guided approach based on the crystal structure of GID1A^9^, targeting residues in the GA-binding pocket predicted to sterically clash with the C-1 side group of sGA. Substitution of Phe27, Tyr31, and Ile126 with smaller residues was expected to enlarge the pocket and facilitate the binding of the synthetic ligands. To identify functional variants, we used a GPS1 biosensor, which uses Förster Resonance Energy Transfer (FRET) to sense binding of GA by the GA receptor module^21^ due to conformational change in GID1C that mediates GA-dependent interaction with the DELLA protein GAI (Fig. 1c). Screening of mutant GPS1 proteins revealed that GPS1 (GID1C^F27G, Y31A, I124A^) showed a slight fluorescence emission ratio changes in response to sGA-1, while GPS1 (GID1C^Y31A^) and GPS1 (GID1C^Y31G^) demonstrated enhanced fluorescence emission ratio changes in response to sGA-1 and sGA-2 (Fig. 1d). All three variant sensors showed no response to GA_3_ and exhibited specific responses to sGAs. The same substitutions were used to construct mGID1 receptors with expanded binding pockets specific to sGA analogs.

To investigate whether these mGID1 variants interact with GAI in an sGA-dependent manner, we performed yeast three-hybrid (Y3H) assays that report GA-dependent interaction of GID1 and GAI. GID1A is considered to be the major GID1 isoform in *Arabidopsis*^11,22^. We introduced the amino acid substitutions identified in the GPS1 screen into GID1A and tested their ability of the three variants to bind GAI in the presence of sGA candidates. This Y3H assay facilitates yeast growth by allowing the production of histidine (His) upon GA-dependent interaction between GID1A and GAI. Yeast expressing wild-type GID1A did not grow in an sGA-1-dependent manner, whereas yeast expressing GID1A^Y31A^ or GID1A^Y31G^ showed robust sGA-1-dependent growth. In contrast, their responsiveness to GA_3_ and GA_4_ was significantly reduced compared to that of wild-type GID1A (Fig. 1e, Extended Data Fig. 2), confirming selective receptor activation by the synthetic ligands. Additionally, yeast co-expressing AtSWEET13, a sugar transporter known to transport GA^23,24^, demonstrated enhanced growth in response to sGA-1 and sGA-2 relative to yeast strains that did not express AtSWEET13 (Fig. 1e), indicating that sGA-1 and sGA-2 are effectively transported by AtSWEET13. The results demonstrate that sGA-1 and sGA-2 function as inert GA analogs selectively recognized by engineered mGID1 variants, providing a modular platform for chemical reconstruction of GA signaling.

### Establishment of the sGA-mGID1 pair in *planta*

To test whether sGA-1 interacts with GID1A^Y31A^ or GID1A^Y31G^ to activate GA signaling in plants, we generated transgenic *Arabidopsis* lines expressing GID1A^Y31A^ or GID1A^Y31G^ fused to GFP under the control of the UBQ10 promoter. As a test system, we used the *ga3ox1 ga3ox2* double mutants, which lack key enzymes in the final step of GA biosynthesis^25^. The mutants exhibit severely reduced endogenous GA levels and a short-root phenotype that can be fully rescued by exogenous GA_3_, providing a genetically sensitized platform to assess synthetic GA signaling in the absence of interference from native hormone activity (Fig. 2a, b). Treatment of *ga3ox1 ga3ox2* plants with sGA-1 alone did not alter root length compared to DMSO controls, confirming that sGA-1 has no activity through endogenous GID1 receptors. In contrast, sGA-1 treatment of either *pUBQ10:GFP-GID1A*^*Y31A*^ *ga3ox1 ga3ox2* or *pUBQ10:GFP-GID1A*^*Y31G*^ *ga3ox1 ga3ox2* lines restored root elongation to levels comparable to GA_3_-treated plants and wild-type controls (Fig. 2a, b, Extended Data Fig. 3). These results demonstrate that sGA-1 specifically activates GA signaling through the engineered receptors and does not interact with native GID1s.

**Fig. 2.**
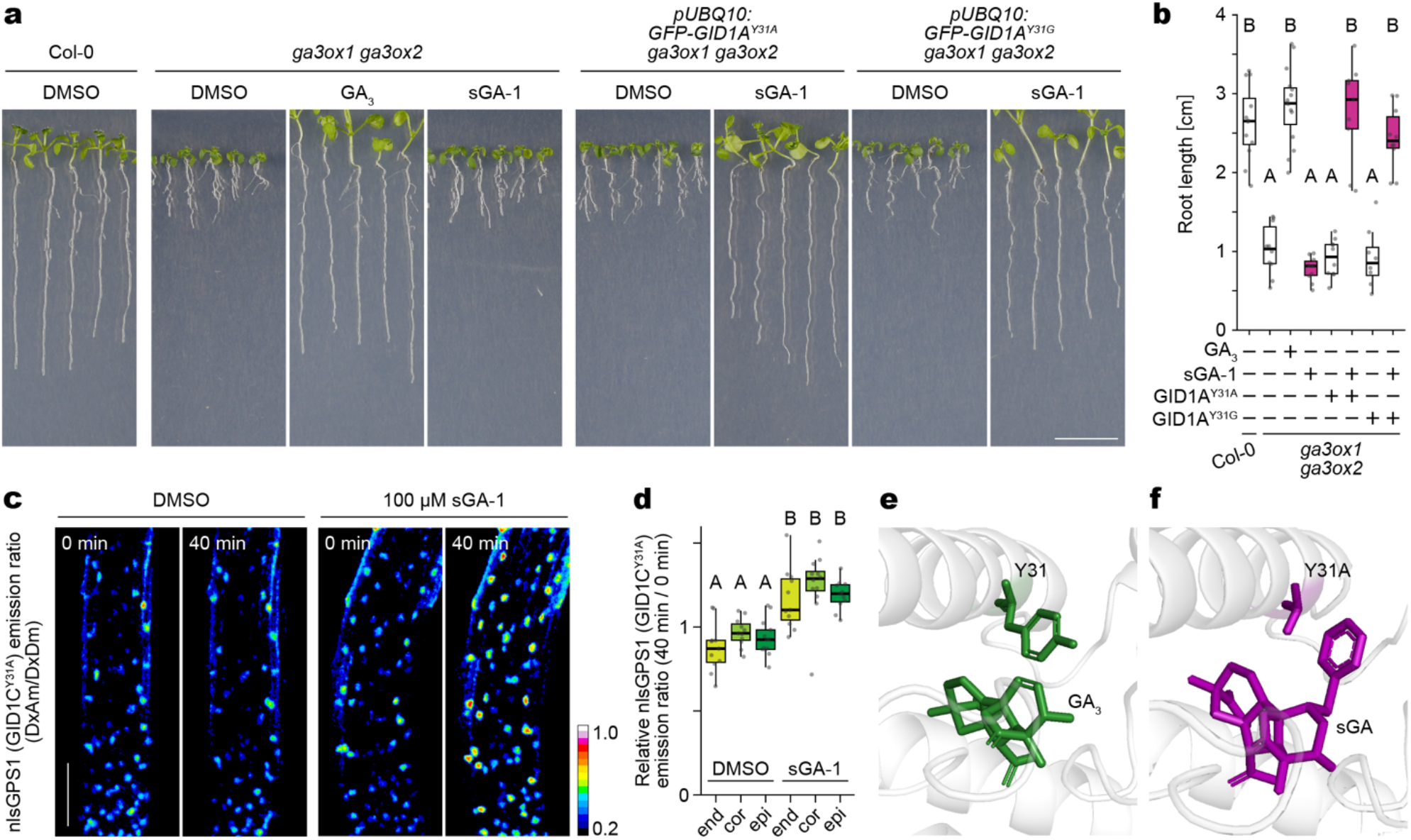
Establishment of sGA-mGID1 system in plants. **a**, Representative images of 9-day-old *Arabidopsis* seedlings grown vertically on agar medium containing DMSO, 1 µM GA_3_, or 1 µM sGA-1. **b**, Total root length of plants in **a** (n ≥ 8 plants). **c**, Longitudinal section of nlsGPS1 (GID1C^Y31A^) emission ratio in an 8-day-old *ga3ox1 ga3ox2* mutant root, showing the region 400–950 µm from the quiescent center (QC). Experiments were repeated three times with comparable results. **d**, Relative nlsGPS1 emission ratio (t = 40 min / t = 0 min) measured in individual nuclei, derived from **c**. The calculation method is shown in Extended Data Fig. 4. The axis labels (end, cor, epi) indicate endodermis, cortex, and epidermis, respectively (n ≥ 11 nuclei from 3 plants). **e**, Crystal structure of GID1A (white) with GA_3_ (green) (PDB 2ZSH). Y31 of GID1A is shown in green. **f**, Predicted docking model of mGID1 (white) with sGA (magenta). The Y31A substitution in mGID1 is shown in magenta. Different letters in **b**,**f** indicate statistically significant differences between samples, as determined by one-way ANOVA analysis with Tukey’s HSD test (p < 0.05). In box plots, a box represents the range from 25^th^ to 75^th^ percentile, the horizontal line marks the median value, and the whiskers capture the extremes within 1.5 times the interquartile range. Scale bars: 1 cm in **a** and 100 µm in **c**.

Since the larger Fluorescein-labeled GA_3_ molecule has been shown to distribute and accumulate in the endodermis^15^, we next examined whether sGA-1 can enter inner root tissues. Using plants expressing a modified GPS1 containing the GID1C^Y31A^ variant, we detected fluorescence ratio changes of the biosensor in response to sGA-1 in the epidermis, cortex, and endodermis (Fig. 2c, d, Supplementary Video 1 and 2; Extended Data Fig. 4), comparable to GPS1 with GA_4_^21^, indicating that sGA-1 diffuses effectively across root tissues and can access multiple layers to activate signaling through mGID1. Based on its inactivity toward native GID1, strong activation of engineered receptors, and efficient tissue permeability, we adopted sGA-1 as sGA ligand and GID1A^Y31A^ as the optimized mGID1 for subsequent experiments (Fig. 2e, f).

### GA signaling restricted to the endodermis is insufficient for proper root development in *Arabidopsis*

GA signaling in the endodermis has long been considered critical for promoting root growth^13,14^, leading to the prevailing assumption that endodermal GA signaling alone might be sufficient to drive root elongation. However, it has remained unclear whether this layer can fully coordinate organ growth in the absence of GA signaling in adjacent tissues. To directly test the specific contribution of the endodermis to GA-mediated root elongation, we expressed *GFP-mGID1* under the control of the *SCARECROW* (*SCR*) promoter^26–28^, which is active specifically in the endodermis, in the *ga3ox1 ga3ox2* GA-deficient background (Fig. 3a). Seedlings were first germinated on sGA-free medium and then transferred to sGA-containing medium before measuring root elongation (Extended Data Fig. 5a). The *pSCR:GFP-mGID1 ga3ox1 ga3ox2* line showed a significant increase in root elongation compared with *ga3ox1 ga3ox2* mutants, indicating that endodermal GA signaling partially promotes elongation (Fig. 3b). However, root growth remained substantially below that of *pUBQ10:GFP-mGID1 ga3ox1 ga3ox2* seedlings, in which mGID1 was produced-and GA signaling was activated-throughout the root, and full elongation was restored (Extended Data Fig. 5b, c; Fig. 2a, b). These results reveal that GA signaling confined to the endodermis is not sufficient for full root growth, indicating the need for coordinated GA activity across multiple cell layers to achieve proper root development.

**Fig. 3.**
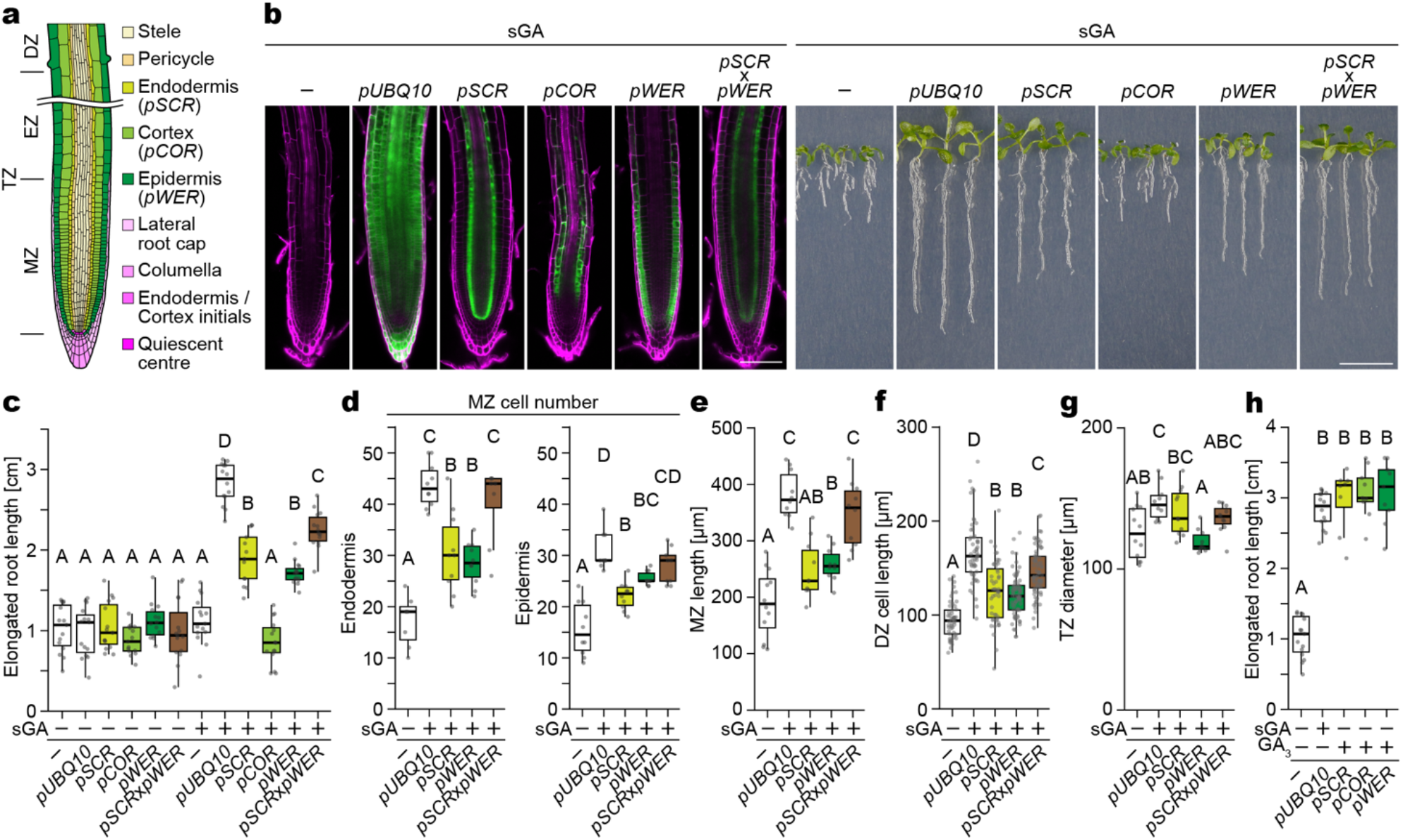
Radial coordination of GA signaling across root tissue layers in *Arabidopsis*. **a**, Schematic diagram of the Arabidopsis root tip showing major tissue layers and growth zone such as meristematic zone (MZ), transition zone (TZ), elongation zone (EZ), and differentiation zone (DZ). **b**, Confocal images of root tips (left half) and corresponding seedling images (right half) from 9-day-old *ga3ox1 ga3ox2* mutants expressing *GFP-mGID1* under cell type-specific promoters or from *ga3ox1 ga3ox2*. Plants were treated with DMSO or 1 µM sGA following the protocol described in Extended Data Fig. 5a; representative images of plants treated with sGA are shown. Promoters driving *GFP-mGID1* expression are indicated above each image. F_1_ seedlings from crosses between *pSCR* and *pWER* lines were used. Magenta indicates PI and green indicates GFP-mGID1. Scale bars, 100 µm (left) and 1 cm (right). **c**, Total elongated root length of plants shown in **b** (n ≥ 13 plants). **d**-**g**, Quantitative phenotypic analysis of the plants in **b**: Number of endodermal (n ≥ 7 plants) or epidermal (n ≥ 9 plants) cells in the MZ (**d**), MZ length (**e**) (n ≥ 10 plants), cell length in the DZ (**f**) (n ≥ 48 cells from 4 plants), and TZ diameter (**g**) (n ≥ 10 plants). Analyses include *ga3ox1 ga3ox2* mutants, lines with whole-root GA signaling (pUBQ10), and cell type-specific activation lines. **h**, Elongated root length of *ga3ox1 ga3ox2* mutants, the lines with whole-root GA signaling, and cell-specific GFP-mGID1 expressing lines treated with GA_3_ (n ≥ 8 plants). GA_3_ treatment was performed as described for sGA in **b**. Different letters in **c**-**h** indicate statistically significant differences between samples, as determined by one-way ANOVA analysis with Tukey’s HSD test (p < 0.05). Exact *P*-values are described in Extended Data. In box plots, box represents the range from 25^th^ to 75^th^ percentile, the horizontal line marks the median value, and the whiskers capture the extremes within 1.5 times the interquartile range.

### Spatial coordination of GA signaling in *Arabidopsis* root development

To investigate the contribution of GA signaling of cell types beyond the endodermis, *GFP-mGID1* was expressed under the cortex-specific *COR* promoter^29^ or the epidermis-specific *WEREWOLF* (*WER*) promoter^30^ (Fig. 3a). If the endodermis acts as the sole driver of elongation, activating GA signaling in the cortex or epidermis would be expected to have minimal effects. Instead, activation of GA signaling solely in the epidermis significantly increased root elongation, whereas cortex-specific signaling did not significantly enhance growth compared to the *ga3ox1 ga3ox2* mutants (Fig. 3b, c), indicating a critical role for epidermal GA responses. Using the *SCR* and *WER* promoters together, concurrent activation in both the endodermis and epidermis resulted in greater root elongation than either tissue layer alone (Fig. 3b, c), indicating that GA signaling in the endodermis and epidermis acts cooperatively to drive root growth. Interestingly, in etiolated hypocotyls, activation of GA signaling in the cortex promoted elongation, whereas application of sGA did not activate signaling when mGID1 was expressed under the *WER* promoter failed (Extended Data Fig. 6a-c). Thus, *WER*-driven activation enables root-specific promotion of elongation.

At the cellular level, organ growth is governed by the interplay between cell proliferation and elongation^31^. To elucidate the cellular mechanisms underlying root elongation, we quantified cell proliferation and elongation in lines exhibiting enhanced root growth. Endodermis-specific GA signaling increased cell division not only in the endodermis but also in the epidermis; likewise, epidermis-specific activation stimulated proliferation in both epidermis and endodermis (Fig. 3d). These findings are consistent with intercellular communication and signaling pathways that propagate proliferative cues across tissue layers^32,33^. Specifically, GA-induced elongation in one layer promotes compensatory division in adjacent tissues, i.e., endodermal elongation induces epidermal division, and vice versa, downstream of GA perception. When GA signaling was simultaneously activated in both the endodermis and epidermis, cell division in both layers reached levels comparable to those observed in roots where GA signaling was induced throughout the entire root using the *UBQ10* promoter, resulting in normal root elongation (Fig. 2a, 3d). Consistently, activation of GA signaling in both the endodermis and epidermis layers restored the length of the meristematic zone (MZ) (Fig. 3e). Measurement of cell elongation in the differential zone (DZ), using root hair spacing as a proxy, revealed cell elongation when GA signaling was active in either the endodermis or epidermis and further enhancement when both layers were activated simultaneously (Fig. 3f). Although root length was comparable between endodermis- or epidermis-specific activation (Fig. 3b, c), the transition zone (TZ) diameter was smaller in epidermis-specific lines compared to endodermis-specific lines (Fig. 3g). Treatment with GA3 restored root elongation in all three lines to the same level as the p*UBQ10* lines, confirming that the partial responses observed with tissue-specific activation were due to restricted signaling domains rather than reduced signaling capacity (Fig. 3h). Cell type-specific activation of GA signaling demonstrates that GA perception and signaling in the endodermis and epidermis cooperatively promotes cell division and elongation, thereby orchestrating and ensuring entire root elongation in *Arabidopsis*.

### Spatial segregation of GA biosynthesis and signaling in *Arabidopsis* roots

A detailed understanding of where GAs are synthesized and perceived is essential for elucidating how local hormone dynamics coordinate root growth. Although GA biosynthesis has been extensively studied at the transcriptional level, the precise cellular localization of GA biosynthetic enzymes in roots remains unclear. Previous analyses using promoter-GUS reporters^25,28,34^ and single-cell RNA sequencing visualized on the Root Cell Atlas website tool (https://rootcellatlas.org)^35–41^ have provided valuable insights into the expression patterns of *GA3ox1* and *GA3ox2*. However, these approaches detect transcriptional activity rather than protein distribution, and the results differ partly in terms of cell-type specificity. To determine GA3ox protein distribution, we generated GA3ox1-GFP and GA3ox2-GFP lines in the *ga3ox1 ga3ox2* mutant background, in which both fusion proteins fully restore root length defects (Extended Data Fig. 7a, b). Both GA3ox1-GFP and GA3ox2-GFP were accumulated in the pericycle of the DZ, while GA3ox2-GFP was additionally detected in the endodermis and cortex in the elongation zone (EZ), as well as the quiescent center (QC) and columella cells, consistent with the previous promoter-GUS analysis^25^ (Fig. 4a, Extended Data Fig. 7c). GA3ox1-GFP showed weak, non-cell-type-specific accumulation in the root tip (Fig. 4a). Notably, GA3ox2-GFP was absent from the epidermis, where GA signaling strongly promotes root elongation, and, although GA3ox2-GFP was present in the cortex, the cortex layer contributed little to GA-induced cell division or elongation under our experimental conditions. These findings reveal a partial spatial separation between GA biosynthetic and GA signaling sites, indicating that bioactive GA_4_ is transported from the cortex, QC, and columella to the epidermis and possibly to the endodermis to coordinate root growth (Fig. 4b, c). This spatial decoupling implies the existence of intercellular GA transport or storage mechanisms that integrate hormone production and signaling across tissue layers, ensuring coordinated and balanced organ growth.

**Fig. 4.**
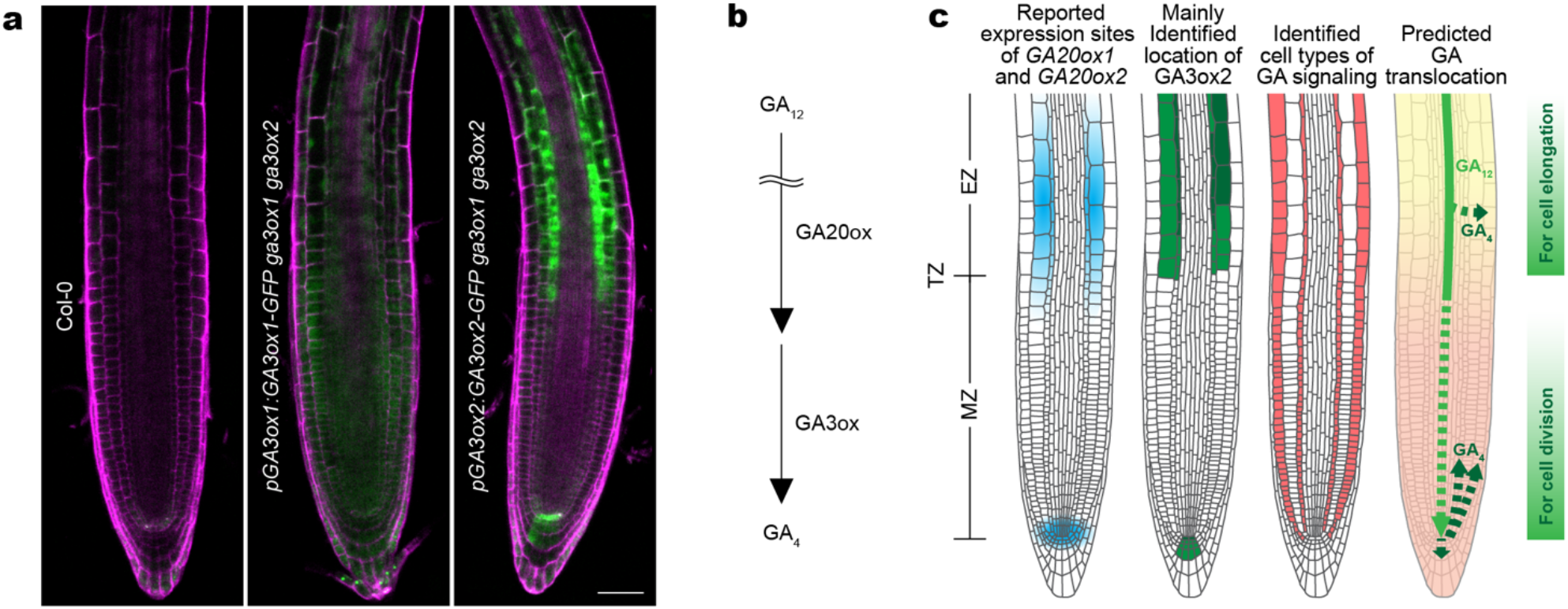
GA biosynthesis, signaling and translocation pathways. **a**, Confocal images of root tips from 5-day-old Col-0, *pGA3ox1:GA3ox1-GFP ga3ox1 ga3ox2*, and *pGA3ox2:GA3ox2-GFP ga3ox1 ga3ox2* seedlings stained with propidium iodide (PI, magenta) and imaged for GFP fluorescence (green). Scale bar, 50 µm. The experiments were repeated three times with similar results. **b**, Simplified diagram of the GA biosynthetic pathway. For clarity, intermediates such as GA_9_ are not shown here. **c**, Conceptual schematic summarizing the identified sites of GA biosynthetic and signaling and the predicted routes of GA translocation. The approximate expression sites of *GA20ox1* and *GA20ox2* are adopted from Barker *et al*. (2021). The dashed arrows indicate the possible presence of GA translocation.

## Discussion

Precise spatial control of hormone signaling is crucial for coordinating organ growth in multicellular plants, yet there are limited tools available for manipulating hormone activity with cellular precision. Here, we establish a synthetic GA-receptor system that enables spatially precise reconstitution of GA signaling in plants, which was designed based on a synthetic approach used to study auxin signaling^42^. The system comprises a biologically inactive GA analog (sGA) and a modified GID1 receptor (mGID1) engineered to selectively bind sGA through an expanded ligand-binding pocket. By expressing mGID1 in defined cell types and applying sGA, we achieved cell type-specific activation in *Arabidopsis thaliana*. These findings redefine the spatial logic of GA signaling and introduce a broadly applicable platform for the precise and synthetic control of plant hormone responses.

Our results revise the prevailing model that GA action in the endodermis alone is sufficient to drive root elongation^13,14^. While endodermis-restricted GA signaling partially restored growth in GA-deficient mutants, full recovery required additional activation in the epidermis. This cooperative requirement highlights functional independence between the two layers and indicates that root elongation depends on coordinated expansion and proliferation across tissues. Activation of GA signaling in one layer enhanced growth in adjacent layers, indicating the presence of this coordination remains unsolved, both chemical signals and biophysical coupling between layers have been proposed to contribute to synchronizing growth during plant morphogenesis^32,33^.

The coordinated requirement of GA signaling in both the endodermis and epidermis underscore their complementary roles in root growth. The endodermis functions as an inner growth regulator, setting the pace of elongation through cell expansion and signaling feedback to surrounding layers^43,44^. In contrast, the epidermis serves as the outer executor of elongation^43,45^, directly responding to environmental and mechanical cues^46,47^. Synchronization of GA responses across these two layers likely ensures balanced expansion and structural integrity of the root. Without such coordination, differential growth between inner and outer tissues could generate physical stress between cell layers, leading to reduced elongation or deformation, as predicted by a simulation model^48^. Thus, our findings indicate that GA acts as a molecular integrator, harmonizing growth between inner and outer tissues to achieve coherent organ elongation.

Our results also offer a mechanistic explanation for elevated GA accumulation observed in the cortex and epidermis^21^. Activation of GA signaling exclusively in the cortex did not promote root elongation, despite the accumulation of GA3ox enzymes in this layer, which indicates that the cortex functions primarily as a source or conduit of bioactive GA rather than a major signaling site. The spatial separation of GA biosynthesis and signaling implies that the cortex may serve as a temporary reservoir for bioactive GA_4_. Interestingly, GA signaling in the cortex is associated with the regulation of legume nodulation in pea^49^, implying that cortical GA responses may have evolved specialized developmental roles distinct from root elongation. Limited DELLA expression and mechanical constraints within the cortex may further restrict ability of this tissue to translate GA activation into organ-level growth.

Beyond the biological insights that the sGA-mGID1 system has already provided, the system represents a versatile platform for synthetic control of hormone signaling. Because sGA is orthogonal to native GA pathways, it allows selective reconstitution of GA signaling in defined cell types, tissues, or developmental stages without perturbing endogenous hormone dynamics. This framework opens opportunities for rational manipulation of plant architecture. For instance, select manipulation could promote root elongation without increasing shoot height, thereby enhancing nutrient uptake and reducing fertilizer dependency. More broadly, pairing synthetic ligands with engineered receptor or degron modules^42,50^ could enable programmable control of diverse hormonal processes, advancing synthetic biology toward spatially resolved reprogramming of plant development.

## Methods

### Synthesis of GA analogs

Details for the synthesis of sGA-1 and sGA-2 are described in the Supplementary Note.

### Screening of modified GID1 using GA biosensor GPS1

The yeast strain BY5625 [*MATα ura3-52 trp1 leu2-*Δ*1 his3-*Δ*200 pep4::HIS3 prb1-*Δ*1*.*6R can1*] was obtained from the National Bio-Resource Project (NBRP, Japan) and used to express gibberellin biosensor GPS1^21^. To generate the plasmids for screening the GID1C variants, namely *pDRf1-GPS1, pDRf1-GPS1 (GID1C*^*F27G, Y31A, I124A*^*), pDRf1-GPS1 (GID1C*^*Y31A*^) and *pDRf1-GPS1 (GID1C*^*Y31G*^), GPS1 was cloned into the pDONR221 vector by Gateway BP Clonase II enzyme (Invitrogen). Site-directed substitutions were introduced by inverse PCR using PrimeSTAR GXL DNA Polymerase (Takara) and NEBuilder HiFi DNA Assembly (NEB). The resulting pDONR221-GSP1 variants were transferred into the pDRf1 vector^51^. Yeast transformation was performed using a lithium acetate/PEG method following a previous study^24^, and transformants were selected on SD (-Ura) medium. Single colonies were incubated overnight in 3 mL YPD at 30 °C, then 400 µL of culture each was transferred to new 3 mL YPD liquid media and incubated for 3 hours at 30 °C. Cells were centrifuged at 2,300 x g for 30 seconds at 4 °C, and washed three times with 1 mL PBS buffer (137 mM NaCl, 2.7 mM KCl, 10 mM Na_2_HPO_4_, 1.8 mM KH_2_PO_4_, pH 7.4, 4 °C). Pellets were frozen in liquid nitrogen for three minutes, mixed with zirconia/silica beads (0.5 mm; BioSpec), and disrupted using a beads shocker homogenizer (25 times/seconds; QIAGEN). Cell lysates were resuspended in 700 µL PBS buffer by vortex and centrifuged at 15,000 x g for 10 minutes at 4 °C. The resulting supernatants were used in fluorescence analysis in clear-bottom microtiter plates using a SPARK Microplate Reader (Tecan). 50 µL of supernatants were mixed with 50 µL PBS buffer containing either 2 µM GA_3_, 2 µM sGA-1, 2 µM sGA-2 or DMSO. Fluorescence emission spectra from 465 to 567 nm (step size 2 nm) were recorded with excitation at 430 nm. PBS was used as a negative control for background subtraction.

### Yeast three-hybrid assay

Yeast three-hybrid (Y3H) assays were performed as previously described^23,24^. The yeast strain PJ69-4A [*MATa gal4*Δ *gal80*Δ *ura3-52 his3-200 leu2-3,112 trp1-901 LYS2::GAL1-HIS3 ade2::GAL2-ADE2 met2::GAL7-lacZ*] was obtained from the National Bio-Resource Project (NBRP, Japan). Yeast cells were transformed with pDEST22-GAI and either pDEST32-GID1A or mutated pDEST32-GID1A and either pDRf1-GW (empty vector control) or pDRf1-AtSWEET13 by lithium acetate/PEG transformation. Site-directed substitutions were introduced by inverse PCR using PrimeSTAR GXL DNA Polymerase (Takara) and NEBuilder HiFi DNA Assembly (NEB). Yeast colonies were incubated in SD (-Leu, -Trp, -Ura) liquid media and incubated overnight at 30°C. Cultures were sequentially adjusted to OD_600_ = 1.0, 0.1, 0.01, and 0.001. Cell suspensions (5 µL) were spotted on SD + His (-Leu, -Trp, -Ura) or selective SD (-Leu, -Trp, -Ura, -His) media containing either 0.1 μM GA_3_, 1 μM GA_3_, 1 nM GA_4_, 10 nM GA_4_, 1 μM sGA-1, 1 μM sGA-2 or DMSO, and incubated for 3 days at 30°C. Plates were photographed. Yeast colonies obtained from three independent transformations were used for three biological replicates.

### Molecular docking

To construct models of mGID1 bound to the sGA-1 synthetic molecule, molecular docking simulations were performed using AutoDock Vina 1.1.2^52^. The three-dimensional (3D) model of mGID1 was generated using Alphafold2^53^. The 3D structure of sGA-1 was constructed, and its geometries were optimized at the Hartree-Fock/6-31G(d) level using Gaussian 16 (Gaussian.com). Prior to molecular docking, PDB files were converted to PDBQT format using OpenBabel 3.1.1^54^. Docking sites were identified by superimposing the apo-mGID1 model. Grid boxes were defined to enclose sGA-1 within mGID1, following the methodology of a previous study^24^. Docking poses for each ligand were ranked based on binding free energy estimations and the population sizes of the resulting clusters. The docking pose with the lowest binding free energy within the most populated cluster was selected as the final conformation.

### Plant materials

The *Arabidopsis ga3ox1 ga3ox2* mutant obtained from the *Arabidopsis* Biological Resource Center (ABRC) was used in this study. To generate *pUBQ10:GFP-GID1A*^*Y31A*^ and *pUBQ10:GFP-GID1A*^*Y31G*^ binary vectors, *GID1A* cDNA was PCR-amplified and cloned into the pDONR221 entry vector by Gateway BP Clonase II (Invitrogen). Site-directed substitutions (Y31A or Y31G) were introduced by inverse PCR using PrimeSTAR GXL DNA Polymerase (Takara) and NEBuilder HiFi DNA Assembly (NEB). *GID1A*^*Y31A*^ and *GID1A*^*Y31G*^ were fused with GFP in pGWB406m^55^ through LR recombination, and the *GFP-GID1A*^*Y31A*^ and *GFP-GID1A*^*Y31G*^ fragments were cloned into a binary vector, which contains pUBQ10 promoter and HSP18.2 terminator, kindly provided by Dr. Akira Yoshinari (Nagoya University, Japan). *pUBQ10:GFP-GID1A*^*Y31A*^ and *pUBQ10:GFP-GID1A*^*Y31G*^ were introduced into *Agrobacterium tumefaciens* strain GV3101:pMP90 to transform *ga3ox1 ga3ox2* mutant plants through the floral dip method^56^.

For cell-specific expression, *pSCR:GFP-mGID1, pCOR:GFP-mGID1*, and *pWER:GFP-mGID1* binary vectors were generated using Gateway technology (Invitrogen) as described previously^57^. A GFP sequence with 3×GGGGS linker^58^ was inserted before the start codon of *GID1A*^*Y31A*^ in pDONR221 by inverse PCR. Promoter entry clones (pSCR-p4p1r, pCOR-p4p1r, and pWER-p4p1r) were generated corresponding to the 2000-bp, a 1256-bp, or a 3729-bp sequence upstream from the start codon of the *SCARECROW* (AT3G54220), AT1G09750, and *WEREWOLF* (AT5G14750), respectively. The three entry clones, promoter, GFP-mGID1, and t35S-p2p3 were transferred into the binary vector pBm43GW^59^. The final constructs were introduced into *Agrobacterium tumefaciens* GV3101:pMP90 to transform *ga3ox1 ga3ox2* mutant. The *pSCR × pWER:GFP-mGID1 ga3ox1 ga3ox2* was obtained by crossing the *pSCR:GFP-mGID1 ga3ox1 ga3ox2* and *pWER:GFP-mGID1 ga3ox1 ga3ox2*, and F_1_ progeny were used for the analysis.

To generate *pGA3ox1:GA3ox1-GFP* and *pGA3ox2:GA3ox2-GFP* binary vectors, genomic DNA regions containing sequences 2709-bp or a 3099-bp upstream from the start codon and sequences 738-bp or a 1036-bp downstream from the stop codon were amplified by PCR and cloned into pGEM-T Easy Vector (Promega), respectively. A GFP sequence with 3×GGGGS was inserted before the stop codon of *GA3ox1* or *GA3ox2*. The resulting constructs were digested by *Not*I and integrated into the pBIN40 binary vector by ligation^59^. The constructs were introduced into the *ga3ox1 ga3ox2* mutant.

For the nuclear sGA sensor, nlsGPS1(GID1C^Y31A^), a sequence encoding the SV40-derived nuclear localization signal LQRKKKRKVGG^61^, was inserted immediately after the initiation codon of the *pDRf1-GPS1 (GID1C*^*Y31A*^) by PCR. The nlsGPS1(GID1C^Y31A^) fragment was cloned into the *Asc*I and *Apa*I sites of the binary vector containing pUBQ10 promoter and the HSP18.2 terminator (as described above). The resulting construct was introduced into the *ga3ox1 ga3ox2* mutant.

### Plant growth conditions

*Arabidopsis* seeds were surface-sterilized with a solution containing 16.7% (v/v) NaClO (Fujifilm Wako) and 0.0167% (v/v) Triton X-100 (Sigma-Aldrich), washed with sterile water three times, and stored in water in the dark at 4°C for 3 days. After stratification, seeds were grown on half-strength Murashige Skoog (MS) medium with 1% (w/v) sucrose and 1% (w/v) agar at 23 °C under a long-day photoperiod (16 h light/8 h dark). For growth in liquid medium, 2 ml of ½ MS liquid medium with 0.5% (w/v) sucrose was used to grow plants in 6-well plates at 23 °C under a long-day photoperiod. For growth of etiolated hypocotyls, seedlings were grown in 2 ml of ½ MS liquid medium in 6-well plates at 23 °C. For the first three days, plants were grown under long-day photoperiod, followed by five days in complete darkness created by wrapping the 6-well plates in aluminum foil. To analyze root length and etiolated hypocotyl length, ½ MS medium or ½ MS liquid medium containing either DMSO, GA_3_, sGA-1, or sGA-2 was used.

### Confocal microscopy

*Arabidopsis* seedlings (5-9 days old) were stained with 10 μg/mL propidium iodide (PI). After staining, samples were observed under an LSM800 confocal microscope (Zeiss) equipped with a 20x dry objective. Images shown in Fig. 4a were acquired using a SpinSR spinning disk confocal microscope (Olympus) with a 20x dry objective. GFP and PI were excited at 488 and 561 nm and detection was set at 500-546 nm and 576-700 nm, respectively. For GPS1 imaging, PI-stained roots were gently pressed with a 1% agar block containing ½ MS medium. After 5 min, 100 μL of either DMSO or 100 μM sGA-1 (dissolved in ½ MS) was applied from the side of the agar block. Confocal images of nlsGPS1(GID1C^Y31A^) were acquired using a STELLARIS 8 confocal microscope (Leica) equipped with a 20x dry objective. The root was imaged using an ROI scan with two tiled fields to cover the region of interest. Excitation was performed at 448 nm, and emission was collected at 460–500 nm (CFP channel), 525–560 nm (YFP channel). PI was excited at 565 nm, and emission was collected at 610–680 nm.

### Image processing and analysis

Longitudinal time-lapse z-stacks (x, y, z, t) of tissues containing nlsGPS1 (GID1C^Y31A^) were processed in Fiji/ImageJ software (ver. 2.14.0/1,54f). Both CFP and YFP channels were smoothed using a 3D Gaussian Blur (sigma = 1.5 pixels in x, y, and z). To prevent division by zero, a constant value of 5 was added to the CFP channel (Process > Math > Add). For each time point, the ratio channel (DxAm/DxDm) was calculated for every z-slice using the Image Calculator [fluorescence intensity of YFP channel ÷ (fluorescence intensity of CFP channel + 5)], and the resulting 3D ratio stacks were collapsed into 2D images by Average or Max Intensity Z-projection. The emission ratio images shown in the figures correspond to the projected data at t = 0 min and t = 40 min. The relative nlsGPS1 (GID1C^Y31A^) emission ratio was calculated by the value at 40 min divided by the value at 0 min, as shown in Extended Data Fig. 4.

## Data Availability

The docking model of mGID1 with sGA has been deposited and will be available at https://www.modelarchive.org/.

## Acknowledgements

This research was supported by the Japan Society for the Promotion of Science (JSPS) through the grants 20K21424 and 23H02473 to M.N., and the grant 22H00360 to W.B.F.; by the Suntory Foundation of Life Sciences through a SUNBOR grant to M.N.; and by the Japan Science and Technology Agency (JST) Exploratory Research for Advanced Technology (ERATO, JPMJER2403) to M.N. The ITbM is supported by the World Premier International Research Center Initiative (WPI), Japan.

## Author contributions

S.H., W.B.F., and M.N. designed the research. R.Y., K.I., and S.H. developed synthetic GAs. Y.Y., Y.I., and M.N. performed experiments. E.M. performed molecular docking simulations. Y.Y., Y.I., W.B.F., and M.N. have analyzed data. Y.Y., W.B.F., and M.N. wrote the manuscript. All authors discussed the results and commented on the manuscript.

